# The skate spiracular organ develops from a unique neurogenic placode, distinct from lateral line placodes

**DOI:** 10.1101/2023.03.17.533203

**Authors:** J. Andrew Gillis, Katharine E. Criswell, Clare V. H. Baker

## Abstract

The spiracular organ is an epithelial pouch or tube lined with mechanosensory hair cells, found embedded in the wall of the spiracle/spiracular chamber (remnant of the first pharyngeal cleft) in many non-teleost jawed fishes. It is innervated via a branch of the anterior lateral line nerve and usually considered a specialised lateral line organ, although it most likely functions as a proprioceptor for jaw movement. It is homologous to the paratympanic organ: a hair cell-lined epithelial pouch embedded in the wall of the middle ear (which evolved from the spiracular chamber) of birds, alligators and *Sphenodon*. A previous fate-mapping study showed that the chicken paratympanic organ and its afferent neurons originate from a molecularly distinct placode immediately dorsal to the geniculate (first epibranchial) placode. Here, DiI fate-mapping in a cartilaginous fish (little skate, *Leucoraja erinacea*) shows that the spiracular organ derives from a previously unrecognised neurogenic placode immediately dorsal to the geniculate placode and molecularly distinct from lateral line placodes. These findings further support the developmental and evolutionary independence of this unique jawed-vertebrate mechanosensory organ from the lateral line system.

**Summary statement:** Fate-mapping in a cartilaginous fish supports the developmental independence from the lateral line system of the mechanosensory spiracular organ, consistent with its homology with the amniote paratympanic organ.

## Introduction

The spiracular organ (SpO) is an epithelial diverticulum lined with mechanosensory hair cells embedded in the wall of the spiracle/spiracular chamber (remnant of the first pharyngeal cleft) in representatives of all groups of extant jawed vertebrates: cartilaginous fishes; non-teleost ray-finned fishes excluding bichirs; and the lobe-finned lungfishes and coelacanth (see e.g., Allis, 1889; Agar, 1906; Norris and Hughes, 1920; Barry and Bennett, 1989; Johnston, 2022). In sharks and skates (elasmobranchs), the SpO is embedded in connective tissue near the articulation between the hyomandibula and the braincase (Barry and Boord, 1984; Barry et al., 1988a; Barry and Bennett, 1989) (Fig. 1, Fig. 2a). Afferent innervation is provided via a branch of the anterior lateral line nerve (Barry and Boord, 1984) and the SpO is usually considered to be a specialised lateral line organ (Barry and Bennett, 1989; Northcutt, 1989). However, SpO afferents project centrally not only to the mechanoreceptive lateral line nucleus and vestibulocerebellum but also, uniquely, to lateral regions of the reticular formation (Barry and Boord, 1984), and they are excited by flexion of the hyomandibula at the cranial joint, which distorts the SpO (Barry et al., 1988b). These findings led to the proposal that the elasmobranch SpO is a proprioceptor for jaw movement (Barry et al., 1988b; Barry and Bennett, 1989). Based on the anatomy of the closed spiracular pouch, the lungfish spiracular organ was also suggested to act as a proprioceptor for jaw and hyoid/opercular movements: it is compressed by the spiracular cartilage when the jaw closes, and extended or compressed when the opercular apparatus moves outwards or inwards, respectively (Bartsch, 1994).

**Figure 1:**
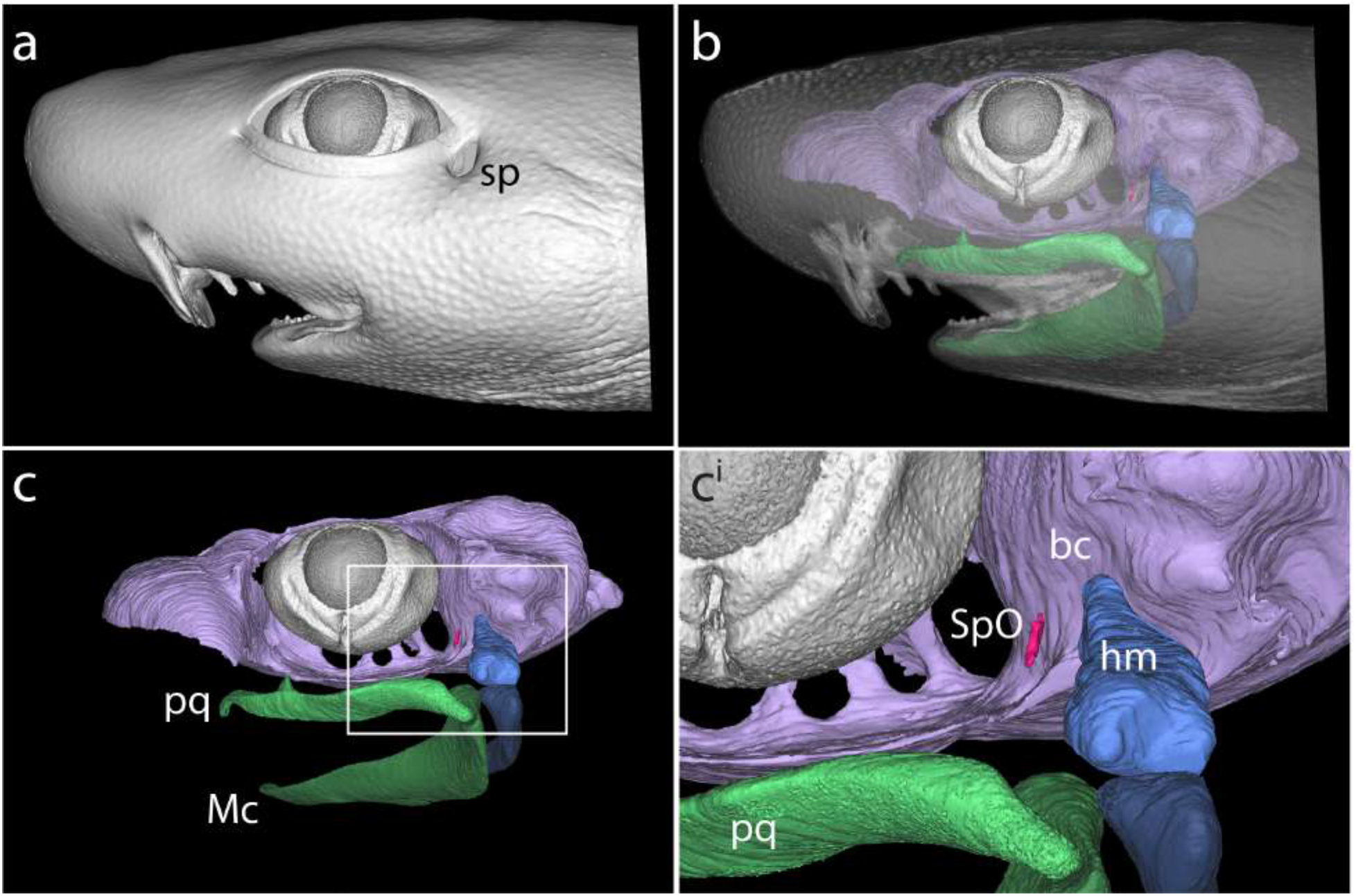
The spiracular organ of cartilaginous fishes. MicroCT reconstruction of the head of a pre-hatchling embryonic shark (*Scyliorhinus canicula*), showing **(a)** the spiracle (*sp*), which sits behind the eye, and which is derived from the first embryonic pharyngeal cleft. **(b**-**c**^**i**^) The spiracular organ (*SpO*, magenta) is an epithelial diverticulum embedded in connective tissue between the hyomandibula (*hm*, blue) and the braincase (*bc*, lilac). The jaws, consisting of Meckel’s cartilage (*Mc*) and the palatoquadrate (*pq*), are highlighted in green.

**Figure 2.**
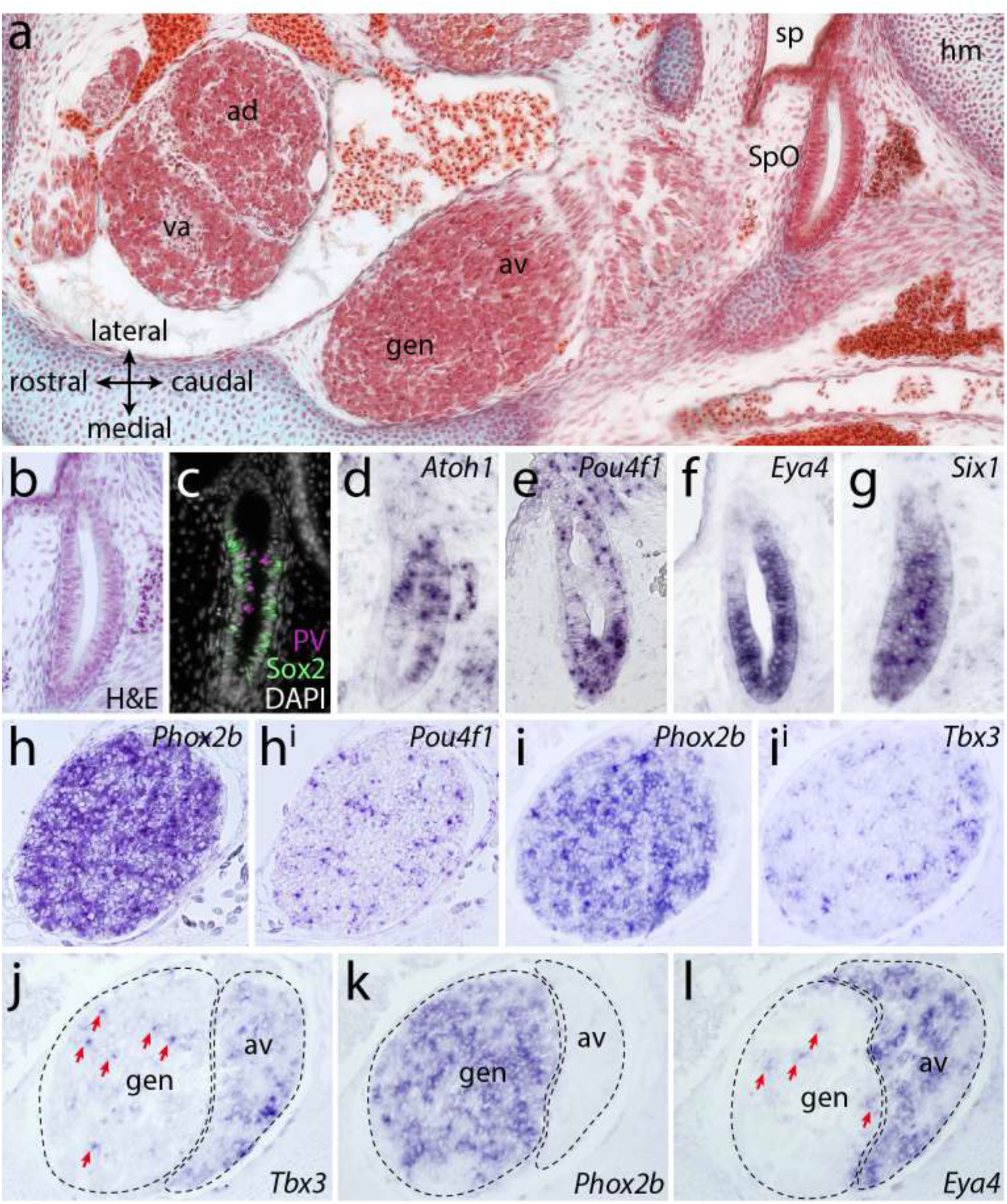
The spiracular organ and nearby cranial sensory ganglia in late-stage skate embryos. **(a)** Masson’s trichrome-stained horizontal (frontal) section through the head of S32 skate embryo at the level of the spiracle, showing the position of the SpO relative to the spiracle and the geniculate ganglion and the vestibuloacoustic ganglion, which form composite ganglia respectively with the anteroventral and anterodorsal lateral line ganglia. **(b)** H&E-stained section of the S32 skate SpO. At this stage, immunofluorescence and *in situ* hybridisation on sections show that the SpO expresses **(c)** the hair cell marker parvalbumin and the supporting cell marker Sox2 in a mutually exclusive pattern, and the transcription factor genes **(d)** *Atoh1*, **(e)** *Pou4f1*, **(f)** *Eya4* and **(g)** *Six1*. (**h-l**^**i**^) *In situ* hybridisation on adjacent horizontal (frontal) paraffin sections through the S32 geniculate ganglion (panels h-i^i^ and j-l are from different embryos; panels j-l are at a more dorsal level than panels h-i^i^ so include the anteroventral lateral line ganglion, which is fused to the geniculate ganglion). Scattered amongst the *Phox2b*-positive neurons in the geniculate ganglion (h,i,k) are putative SpO afferent neurons expressing *Pou4f1* (h^i^), *Tbx3* (i^i^,j) and *Eya4* (l). *Tbx3* and *Eya4* are expressed throughout the anteroventral lateral line ganglion (h,l). *ad*, anterodorsal lateral line ganglion; *av*, anteroventral lateral line ganglion, *gen*, geniculate ganglion; H&E, haematoxylin and eosin staining; *hm*, hyomandibula; *sp*, spiracle; *SpO*, spiracular organ; *va*, vestibuloacoustic ganglion.

In tetrapods, the spiracular chamber evolved into the middle ear cavity and the hyomandibula evolved into the columella/stapes (Clack, 2002). Birds, alligators and the tuatara (*Sphenodon*) possess a mechanosensory hair cell-containing epithelial pouch, the ‘paratympanic organ’ (PTO), embedded in connective tissue in the wall of the middle ear (Simonetta, 1953; Werner, 1963; Neeser and von Bartheld, 2002; von Bartheld and Giannessi, 2011; Giannessi et al., 2013). Given their similar anatomy and shared association with derivatives of the first pharyngeal cleft (i.e., spiracle, middle ear), as well as PTO-afferent projections to vestibular nuclei and the ventral cerebellum (von Bartheld, 1990), it had long been proposed that the SpO and PTO are homologous (see von Bartheld and Giannessi, 2011; Giannessi et al., 2013). However, at odds with the hypothesis of homology was the apparent embryonic origin of the PTO from the geniculate placode, i.e., the first of the ‘epibranchial placodes’ that form dorsocaudal to each pharyngeal cleft and give rise to viscerosensory neurons in the distal sensory ganglia of cranial nerves VII, IX and X (the geniculate, glossopharyngeal and nodose ganglia, respectively) (see Ladher et al., 2010). Ablation and fate-mapping experiments in chicken embryos showed that ectoderm in the region of the geniculate placode forms the PTO as well as geniculate neurons (Yntema, 1944; D’Amico-Martel and Noden, 1983). A geniculate placode origin was also consistent with nerve-tracing experiments that identified PTO-afferent neurons within the geniculate ganglion and collected in a nearby ‘paratympanic extension’ of the geniculate ganglion (von Bartheld, 1990). Clonal analysis via retroviral lineage-labelling yielded similar results (Satoh and Fekete, 2005).

The conflict between an apparent geniculate placode origin for the PTO and its proposed homology with the spiracular organ was resolved by fate-mapping and other experiments demonstrating that the chicken PTO and its afferent neurons arise from a molecularly distinct placode located immediately dorsal to the geniculate placode (O’Neill et al., 2012). Here, we sought to determine experimentally the embryonic origin of the SpO in a cartilaginous fish, the little skate (*Leucoraja erinacea*).

## Results and Discussion

The SpO in the little skate is easily identified on histological sections at stage (S)32, adjacent to the spiracle and near the hyomandibula (Fig. 2a,b). Just as for the chicken PTO (O’Neill et al., 2012), immunofluorescence reveals mutually exclusive expression of a hair cell marker (parvalbumin, a calcium-buffering protein) and the supporting cell marker Sox2 (a SoxB1-class transcription factor) (Fig. 2c). *In situ* hybridisation shows that the SpO expresses *Atoh1* (Fig. 2d), which is essential for hair cell development in bony vertebrates (Bermingham et al., 1999; Millimaki et al., 2007). *Pou4f1* (*Brn3a*) is also expressed in the SpO, primarily concentrated at the distal pole (Fig. 2e). These *Pou4f1*-expressing cells could represent neuroblasts prior to delamination, as seen in the chicken PTO (O’Neill et al., 2012). *Eya4*, a conserved marker for otic and lateral line placodes and their derivatives across jawed vertebrates (O’Neill et al., 2007; Modrell et al., 2011a; Modrell and Baker, 2012), is expressed more uniformly in the SpO (Fig. 2f), as is *Six1* (Fig. 2g), which is expressed by all neurogenic placodes and their derivatives (see Schlosser, 2014; Moody and LaMantia, 2015).

In chicken embryos, the cell bodies of PTO-afferent neurons are scattered within the geniculate ganglion and collected in a small separate ganglion, the ‘PTO ganglion’ (von Bartheld, 1990; O’Neill et al., 2012). PTO neurons are smaller than geniculate neurons and express the vestibuloacoustic neuron marker Pou4f1, but not the epibranchial placode-derived neuron marker Phox2b (O’Neill et al., 2012). Pou4f1-positive neuroblasts delaminate from the PTO placode and, later, from the PTO epithelium (O’Neill et al., 2012). Both *Pou4f1* orthologs are expressed in embryonic zebrafish lateral line ganglia (Kitambi and Chandrasekar, 2021). In the little skate at S32, *in situ* hybridisation on adjacent sections for *Phox2b* versus either *Pou4f1* or the lateral line/vestibuloacoustic ganglion markers *Tbx3* or *Eya4* (O’Neill et al., 2007), revealed a few *Pou4f1*-, *Tbx3*- and *Eya4*-positive cells scattered amongst the *Phox2b*-positive geniculate neurons, as well as throughout the anteroventral lateral line ganglion that is fused with the geniculate ganglion (Fig. 2h-l). Given the presence of Pou4f1-positive PTO-afferent neurons within the embryonic chicken geniculate ganglion (O’Neill et al., 2012), we speculate that the *Pou4f1*-, *Eya4*- and *Tbx3*-positive cells scattered within the skate geniculate ganglion could represent SpO-afferent neurons.

We then attempted to identify a putative SpO placode using candidate gene expression and histology. Classical histological studies in embryonic lungfish (Agar, 1906), gars (Landacre and Conger, 1913; Hammarberg, 1937) and shark (Holmgren, 1940) all suggested that the SpO derives from a primordium that is separate from the lateral line placodes. The PTO placode can be recognised externally in chicken embryos at stage 18 (Hamburger and Hamilton, 1951) as a patch of *Sox2*-positive ectoderm immediately dorsal to the *Pax2*-positive, *Sox3*-positive, *Sox2*-negative geniculate placode, itself lying dorsocaudal to the first pharyngeal cleft (O’Neill et al., 2012). In the little skate at S24, *Pax2* expression identifies the maturing epibranchial placodes caudal to the dorsal region of each pharyngeal cleft (Fig. 3a), as previously reported in shark (O’Neill et al., 2007). The epibranchial placodes also express *Sox3*, in a broader stripe extending further ventrally than *Pax2* at this stage (Fig. 3b). Contrary to our expectations from bony vertebrates, in which *Sox3* is strongly expressed by lateral line as well as epibranchial placodes (e.g., Schlosser and Ahrens, 2004; Modrell et al., 2011b), *Sox3* was not expressed by elongating lateral line primordia in skate (Fig. 3b): these express *Eya4* (Fig. 3c), as expected from shark (O’Neill et al., 2007), and *Sox2* (Fig. 3d). Higher-power views of the first pharyngeal (spiracular) cleft region (insets in Fig. 3a-d) reveal a domain of *Eya4*-positive ectoderm immediately dorsal to the geniculate placode and extending slightly rostrally to it, i.e., in a similar position to the chicken PTO placode (O’Neill et al., 2012), but lacking *Sox2* expression, in contrast to lateral line primordia. The absence of *Sox2* also contrasts with the chicken PTO placode, for which *Sox2* is an early marker, maintained throughout PTO development (O’Neill et al., 2012). Skate embryos develop slowly and it is possible that we missed transient *Sox2* expression. However, we examined multiple embryos and could not detect *Sox2* expression here (although when SpO hair cells form, Sox2 expression is seen in the epithelium; Fig. 2c.)

**Figure 3:**
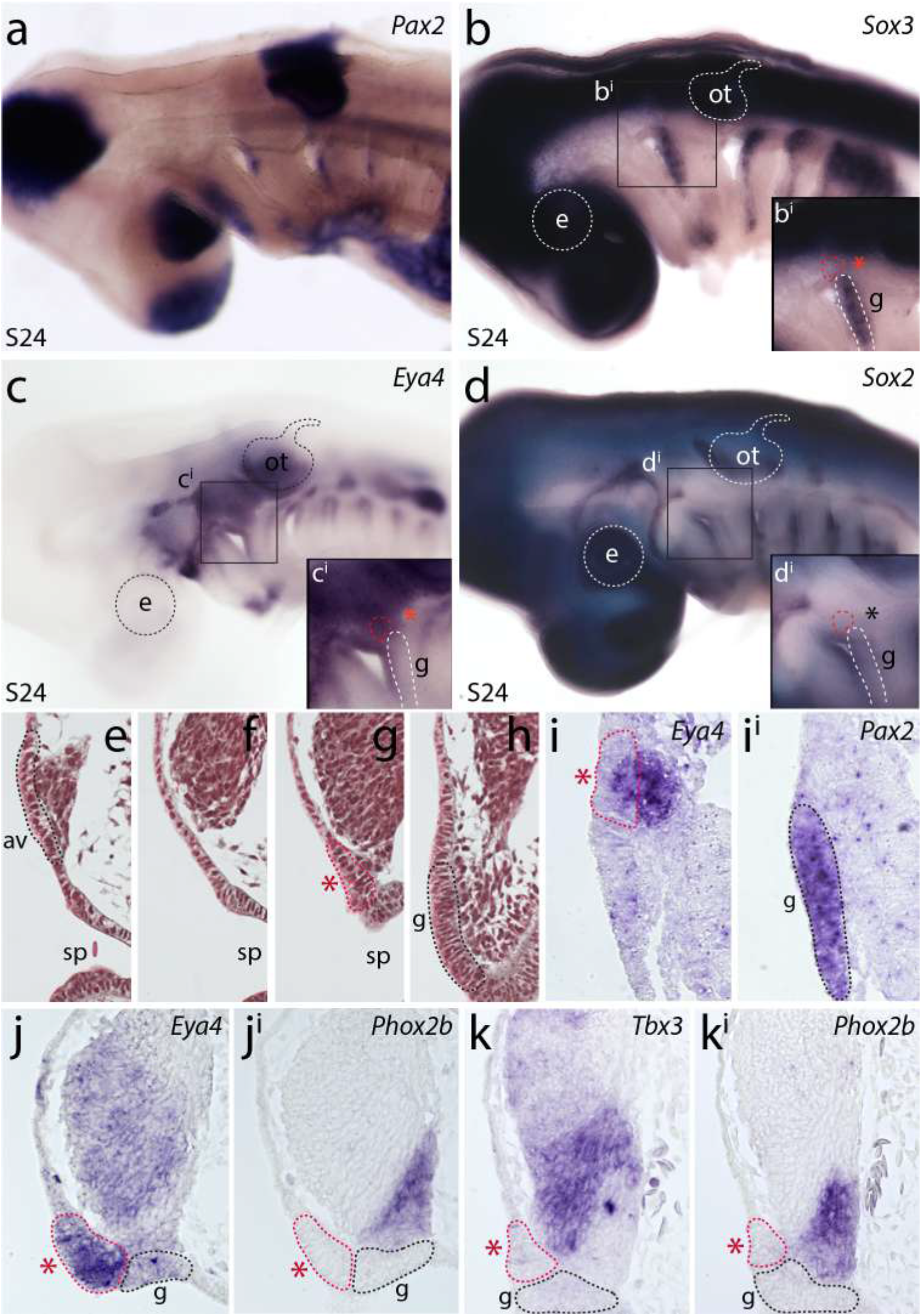
A putative spiracular organ placode in the skate, immediately dorsal to the geniculate placode. Given homology of the amniote PTO and the SpO, the putative SpO placode in skate is expected to lie immediately dorsal to the geniculate (first epibranchial) placode, above the first pharyngeal (spiracular) cleft. **(a-d)** Wholemount *in situ* hybridisation at S24 for *Pax2* (a) and *Sox3* (b) identifies the epibranchial placodes dorsocaudal to each pharyngeal cleft. Ectoderm immediately dorsal to the geniculate placode expresses *Eya4* (c) and is distinct from *Sox2*-positive elongating lateral line primordia (d). Insets show higher-power views of the first pharyngeal (spiracular) cleft region. **(e-h)** Selected transverse histological sections at S24 in a rostral-to-caudal sequence, showing successively a dorsal neurogenic placode with emigrating neuroblasts, expected to be the anteroventral lateral line placode (e); its disappearance (f); then the putative SpO placode (red asterisk) with emigrating neuroblasts (g), which lies immediately dorsal to the geniculate placode with emigrating neuroblasts (h). All these regions of neurogenic placodal ectoderm contribute neuroblasts to the same ganglion. The complete series of histological sections is provided as Fig. S1. **(i**,**i**^**i**^) *In situ* hybridisation on adjacent transverse paraffin sections at S24 reveals a domain of *Eya4*-positive placodal ectoderm (red asterisk) and subjacent mesenchyme, presumably neuroblasts, immediately dorsal to the *Pax2*-positive geniculate placode (i,i^i^). **(j-k**^**i**^) In adjacent transverse sections from a different S24 embryo, *Eya4*-positive placodal ectoderm is seen immediately dorsal to the geniculate placode, here identified as a placodal source of *Phox2b*-positive neuroblasts in the ventral region of the adjacent ganglion (j,j^i^). This domain of placodal ectoderm (red asterisk) is also a source of *Tbx3*-positive neuroblasts, which emigrate immediately dorsal to *Phox2b*-positive geniculate placode-derived neuroblasts (k,k^i^). *ad*, anterodorsal lateral line placode; *av*, anteroventral lateral line placode; *e*, eye; *g*, geniculate placode; *m*, middle lateral line placode; *n1-3*, nodose placodes; ot, otic vesicle; *p*, posterior lateral line placode; *pt*, petrosal placode; *sp*, spiracle; *st*, supratemporal lateral line placode.

Serial rostral-to-caudal histology at S24 confirmed the existence of a domain of neurogenic placodal ectoderm extending dorsally from the geniculate placode above the first pharyngeal (spiracular) cleft, separate from a discrete neurogenic placode lying further rostral and dorsal, all contributing neuroblasts to the same ganglion (Fig. 3e-h; Fig. S1). *In situ* hybridisation on adjacent transverse sections at the level of the second pharyngeal (hyoid) arch showed that the *Eya4* expression domain seen in whole-mount (Fig. 3c) includes placodal (thickened) ectoderm and subjacent mesenchyme (Fig. 3i,i^i^) lying immediately dorsal to the *Eya4*-negative, *Pax2*-positive geniculate placode (Fig. 3i-j), which generates *Phox2b*-positive neurons in the ventralmost wedge of the large adjacent ganglion (Fig. 3j^i^; compare with Fig. 3h). The *Eya4*-positive putative SpO placode also seems to be the source of a tranche of strongly *Tbx3*-positive cells in the ganglion (Fig. 3k), immediately dorsal to the *Phox2b*-positive neuroblasts emigrating from the geniculate placode (Fig. 3k^i^; compare with Fig. 3g). The strongly *Tbx3*-positive cells medially and the weakly *Tbx3*-positive cells dorsally in the composite ganglion also express *Eya4* (compare Fig. 3j,k). The source of the dorsal neuroblasts must be the discrete neurogenic placode identified by serial histology further rostral and dorsal to the putative SpO placode (Fig. 3e). This is presumably the anteroventral lateral line placode, as the mature composite ganglion comprises the geniculate and anteroventral lateral line ganglia (Fig. 2a,j-l; also see Landacre, 1916; O’Neill et al., 2007). Overall, these molecular and histological data suggest that the putative SpO placode (*Eya4*-positive, *Sox2*-negative) is both spatially and molecularly distinct from lateral line primordia (*Eya4*-positive, *Sox2*-positive).

To fate-map the skate SpO directly, we focally labelled ectodermal domains in the dorsal region of the second (hyoid) pharyngeal arch with the lipophilic dye CM-DiI at S24. At S32, when the SpO and cranial sensory ganglia are fully developed (Fig. 2a), embryos were analysed histologically for the presence and distribution of CM-DiI-labelled cells in the SpO and cranial sensory ganglia. In all embryos in which the geniculate placode was targeted at S24 (Fig. 4a,b), CM-DiI-labelled cells were recovered only in the geniculate ganglion at S32, not in the SpO (n=6/6; Fig. 4c-d^i^). In all embryos in which the putative SpO placode was targeted at S24 (Fig. 4e,f), abundant CM-DiI-labelled cells were recovered throughout the SpO, as well as in the geniculate and anteroventral lateral line ganglia (n=12/12; Fig. 4g-h^i^). In six of the twelve SpO placode-targeted embryos, DiI-positive cells were also present in the vestibuloacoustic and anterodorsal lateral line ganglia (not shown). These experiments resolve the embryonic origin of the skate SpO from a domain of placodal ectoderm that is anatomically equivalent to the avian PTO placode (i.e., lies immediately dorsal to the geniculate placode). Our findings also suggest that SpO afferents are broadly distributed across the acousticofacial ganglionic complex (the vestibuloacoustic ganglion contacts the anteroventral lateral line ganglion; see Landacre, 1916; O’Neill et al., 2007).

**Figure 4:**
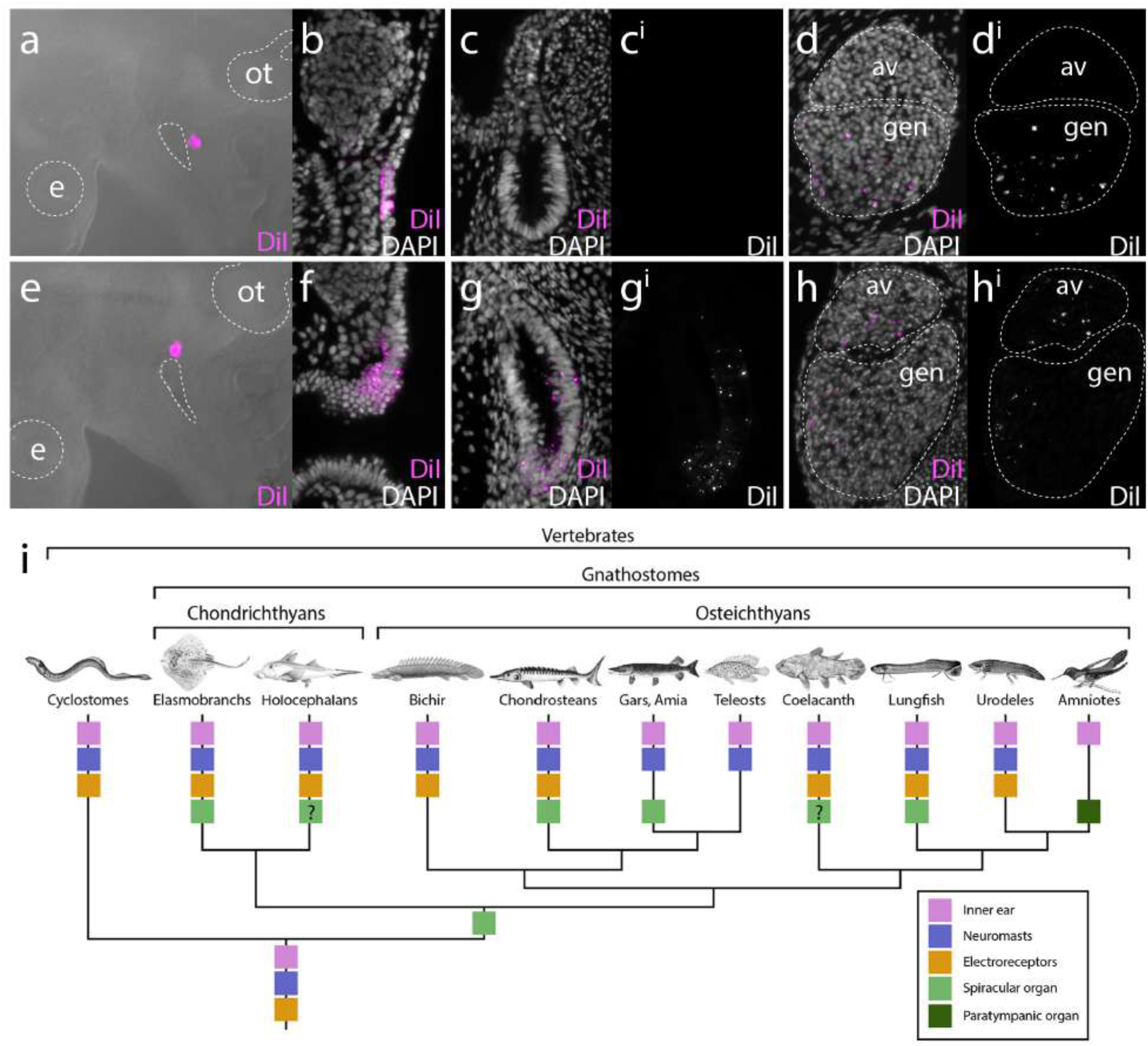
Embryonic origin of the spiracular organ of the skate. **(a-d**^**i**^) Labelling of the geniculate placode with CM-DiI at S24 (a,b) does not label the spiracular organ at S32 (c,c^i^), but abundant CM-DiI-positive neurons are seen in the geniculate ganglion (d,d^i^). **(e-h**^**i**^) Labelling of placodal ectoderm dorsal to the geniculate placode (i.e. the putative SpO placode) with CM-DiI at S24 (e,f) results in abundant CM-DiI-positive cells within the spiracular organ (g,g^i^), as well as CM-DiI-positive neurons within the geniculate and anteroventral lateral line ganglia (h,h^i^). **(i)** Phylogenetic distribution of sensory hair cell-containing sense organs in vertebrates points to an origin of the spiracular organ along the jawed vertebrate stem and its independence from the lateral line system.

Anatomical conservation of the neurogenic SpO/PTO placode in cartilaginous fishes and amniotes further supports the homology of the SpO and PTO (see von Bartheld and Giannessi, 2011; O’Neill et al., 2012), confirming that the last common ancestor of jawed vertebrates possessed a SpO (Fig. 4i). Although the chicken PTO placode expresses *Sox2* (O’Neill et al., 2012), and we were unable to detect *Sox2* expression in the skate SpO placode (only later, in the SpO epithelium), we also report here that elongating lateral line primordia in skate are *Sox3*-negative, in contrast to bony vertebrates (e.g., Schlosser and Ahrens, 2004; Modrell et al., 2011b). These molecular differences between cartilaginous and bony vertebrate placodes are unexpected, but do not call their homology into question.

The ancestral SpO likely had a proprioceptive function for jaw movement, as proposed for elasmobranchs and lungfishes (see Barry and Bennett, 1989; Bartsch, 1994). The molecular distinction between the *Sox2*-negative SpO placode and *Sox2*-positive lateral line primordia in skate suggests at least some degree of developmental independence from the lateral line system. Indeed, the SpO was lost during the evolution of numerous jawed anamniote lineages that retained the mechanosensory lateral line system, including bichirs and teleosts within the ray-finned fishes, and amphibians (Fig. 4i; also see von Bartheld and Giannessi, 2011; O’Neill et al., 2012). The combination of a unique sensory function relating to jaw movement, together with molecular drift from lateral line placode development, may have enabled this evolutionary independence.

Nevertheless, the SpO most likely evolved in the lineage leading to jawed vertebrates via the modification of an existing mechanosensory lateral line organ associated with the first pharyngeal cleft. Pehrson (1949) reported that the spiracular organ in embryonic lungfishes is connected by a “strand” of epithelium to a “neuromast primordium” (also termed “vestigial organ”) in the epidermis, which he described as the most rostral primordium of a transient ‘spiracular line’ (or ‘suprabranchial line’) dorsal to each pharyngeal cleft that disappears during development (Pehrson, 1949). A clear distinction is made between the spiracular organ itself and the “vestigial organs” (neuromast primordia) of the spiracular/suprabranchial line (Pehrson, 1949). As noted by Pehrson (1949) (also see Northcutt, 1989), a ‘suprabranchial line’ of neuromasts has not been reported in any other jawed vertebrate. However, such a line is found in lampreys (Holmgren, 1942; Gelman et al., 2008) (also see Northcutt, 1989). Therefore, it seems plausible that during jaw evolution, selection for responsiveness to jaw movements might have resulted in the modification of a ‘suprabranchial’ lateral line organ associated with the first pharyngeal cleft and its afferents, with developmental drift away from the lateral line placodes. Overall, the phylogenetic distribution of lateral line organs (electrosensory organs as well as neuromasts) and the SpO/PTO, and their impressive morphological variation across taxa, highlights the extent to which vertebrates have repeatedly modified an ancestral sensory repertoire to meet the diverse sensory challenges of their environments.

## Materials and Methods

### Embryo collection and DiI-labelling

*L. erinacea* eggs were obtained from the Marine Biological Laboratory (MBL, Woods Hole, MA, USA) and maintained in a flow-through seawater system at 18ºC to approximately stage (S)24 (Ballard et al., 1993; Maxwell et al., 2008). All animal work complied with protocols approved by the Institutional Animal Care and Use Committee at the MBL. For manipulation, a window was cut in the eggcase and the embryo removed. Embryos were anaesthetised in a solution of buffered ethyl 3-aminobenzoate methanesulfonate salt (100 mg/L; MS-222, Sigma) in seawater. Cell Tracker-CM-DiI (ThermoFisher), diluted 1:10 in 0.3M sucrose from a 5 µg/µl stock in ethanol, was focally injected into ectoderm using a pulled glass capillary needle and a Picospritzer pressure injector. Embryos were allowed to recover in fresh seawater, then replaced in their eggcases and left to develop until S32.

### Histology, immunohistochemistry and *in situ* hybridisation

Following euthanasia by MS-222 overdose (1 g/L), *L. erinacea* embryos were fixed in 4% paraformaldehyde in phosphate-buffered saline (PBS) overnight at 4ºC, rinsed three times in PBS and stored at 4ºC in PBS with 0.01% sodium azide. For histological analysis, embryos were embedded in paraffin as described by Hirschberger and Gillis (2022) and sectioned at 8 µm. Masson’s trichrome staining was performed using the modified protocol described by Witten and Hall (2003). Immunofluorescence with anti-parvalbumin (Merck Millipore MAB1572, mouse IgG1. 1:100) and anti-Sox2 (Abcam ab92494, rabbit monoclonal, 1:200) was performed as described by Hirschberger and Gillis (2022). Nuclei were counterstained with DAPI. *In situ* hybridisation was performed in wholemount and on sections according to the protocol of O’Neill et al. (2012), with modifications according to Gillis et al. (2012). Wholemount *in situ* hybridisation was performed as described by Hirschberger et al. (2021). All sequence data are accessible through NCBI under the following accession numbers: *Atoh1* (OP429207), *Eya4* (JQ425114.1), *Pax2* (OP429214), *Phox2b* (OP429208), *Pou4f1* (OP429209), *Six1* (OP429210), *Sox2* (OP429211), *Sox3* (OP429212), *Tbx3* (OP429213).

## Acknowledgments

The authors thank Prof. Richard Behringer, Prof. Alejandro Sánchez Alvarado, Prof. David Bodznick and the staff of the MBL Marine Resources Center for laboratory and animal support, and Louise Bertrand (Leica Microsystems) for microscopy support.

## Competing interests

The authors declare no competing or financial interests.

## Author contributions

Conceptualization: J.A.G. and C.V.H.B; Methodology: J.A.G.; Formal analysis: J.A.G.; Investigation: J.A.G., K.E.C.; Writing - original draft: J.A.G.; Writing - review & editing: C.V.H.B, J.A.G., K.E.C.; Funding acquisition: J.A.G.

## Funding

This research was funded by the following fellowships and grants to J.A.G.: Royal Society University Research Fellowship UF130182; Plum Foundation John E. Dowling and Laura and Arthur Colwin Endowed Summer Research Fellowships at the Marine Biological Laboratory; and University of Cambridge Isaac Newton Trust grant 14.23z. *Rights Retention Statement:* For the purpose of open access, the author has applied a Creative Commons Attribution (CC BY) licence to any Author Accepted Manuscript version arising from this submission.

---

**Figure S1:**
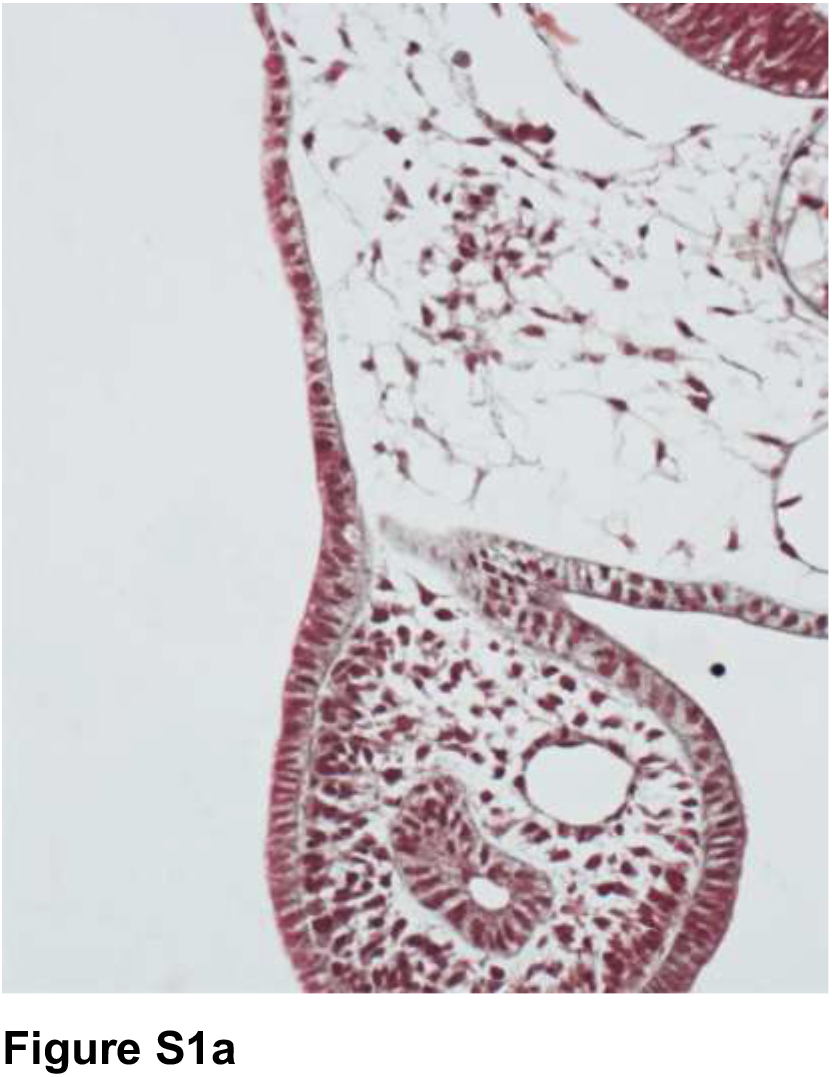
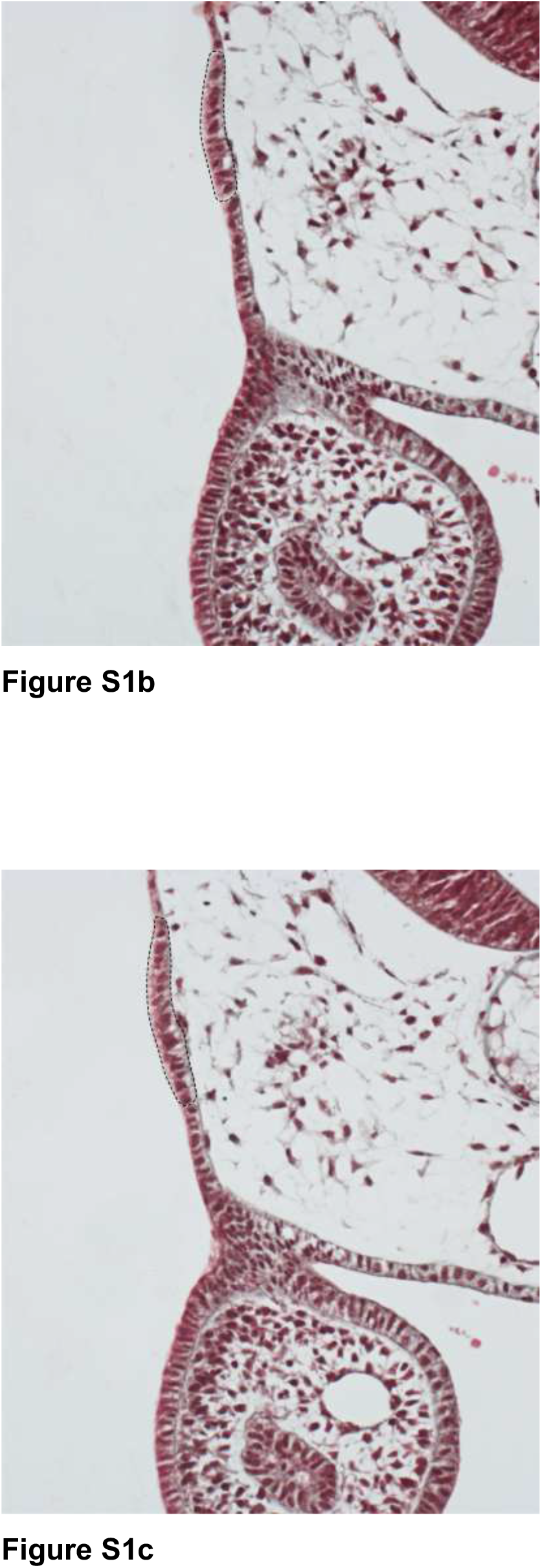
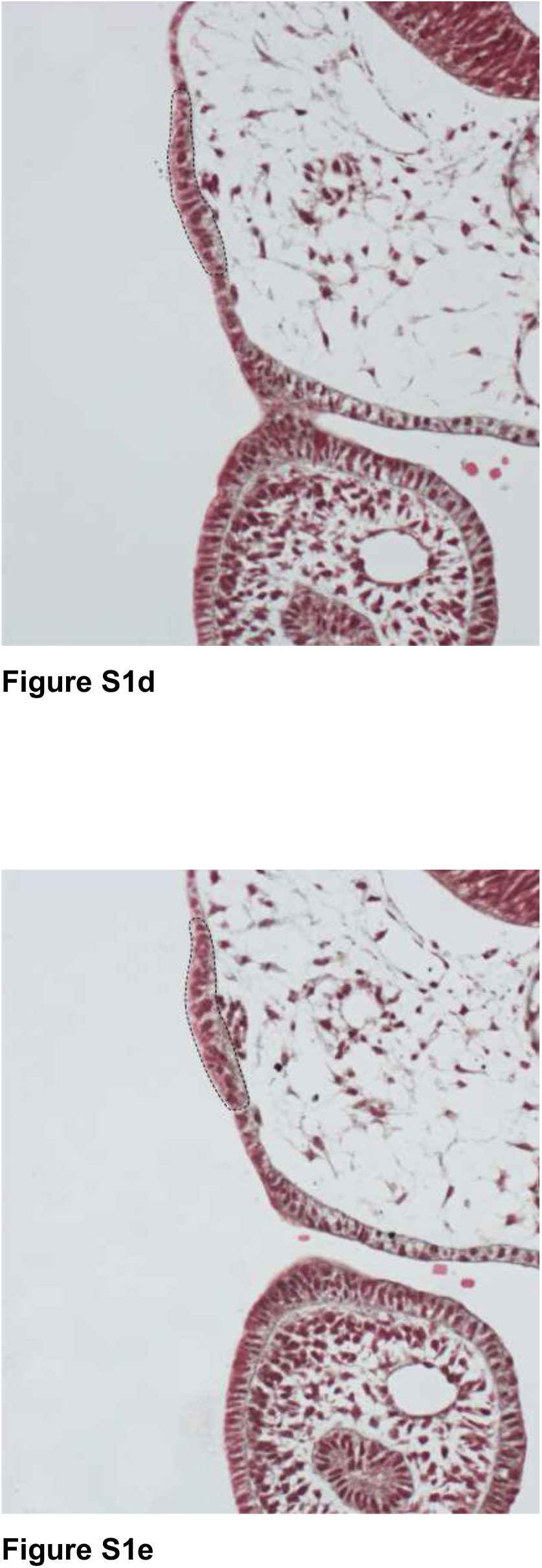
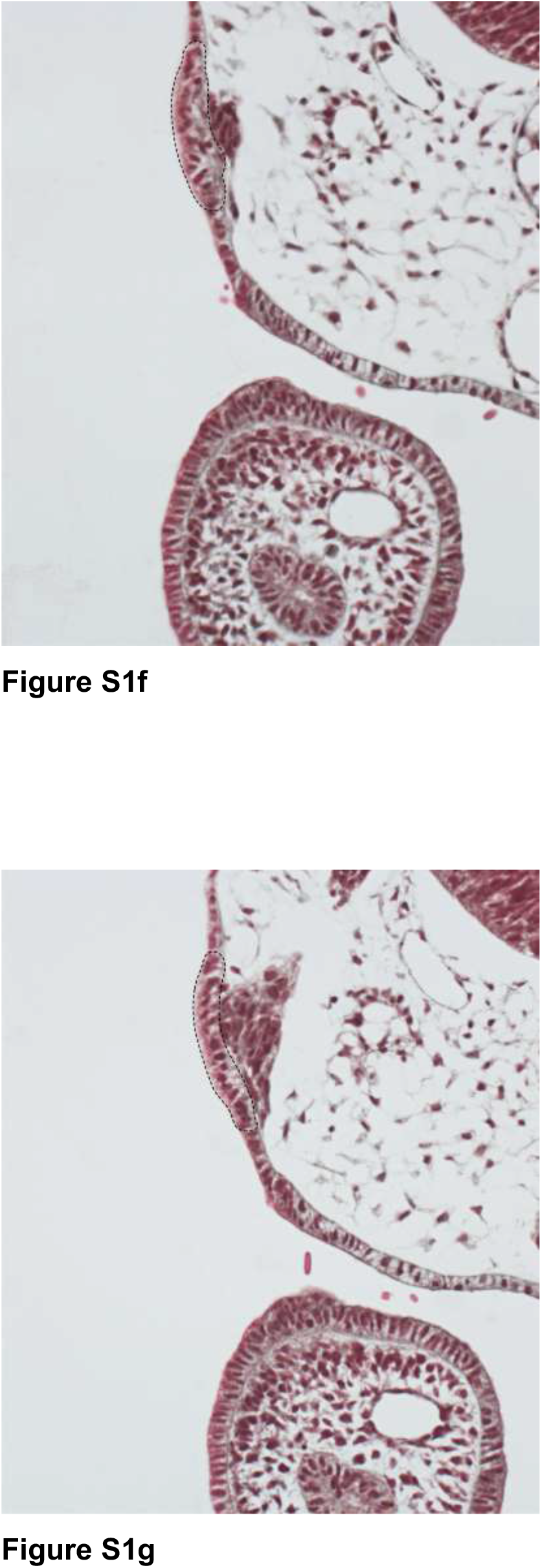
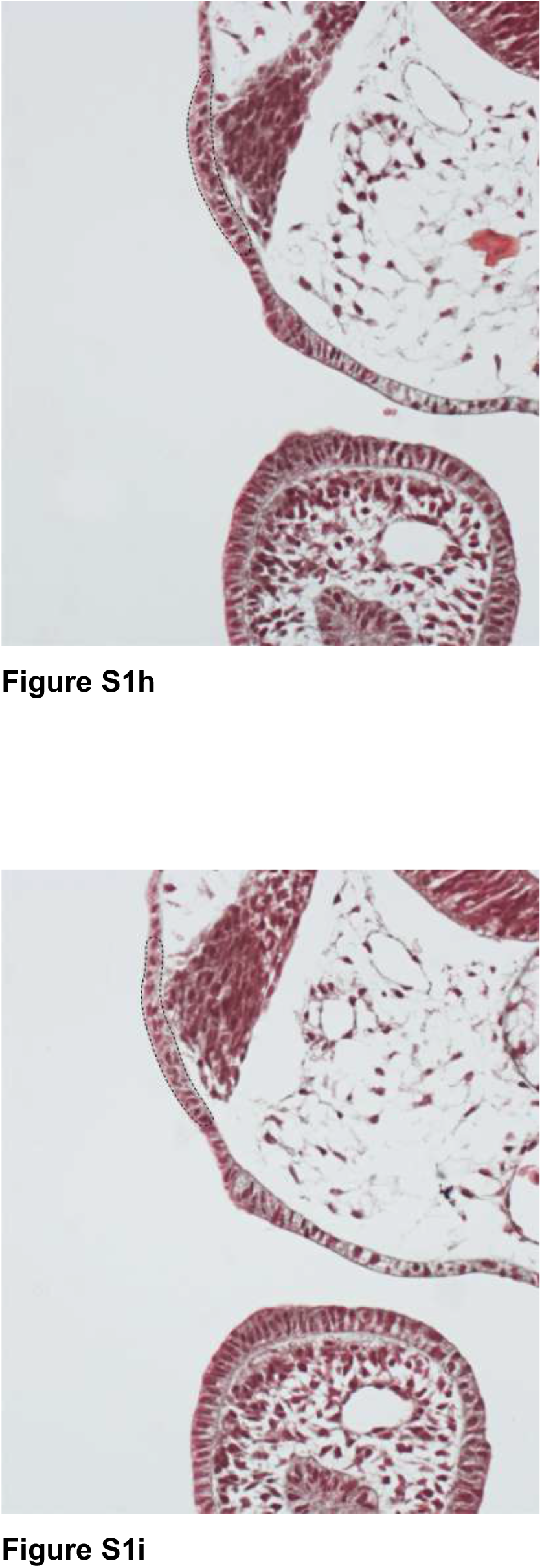
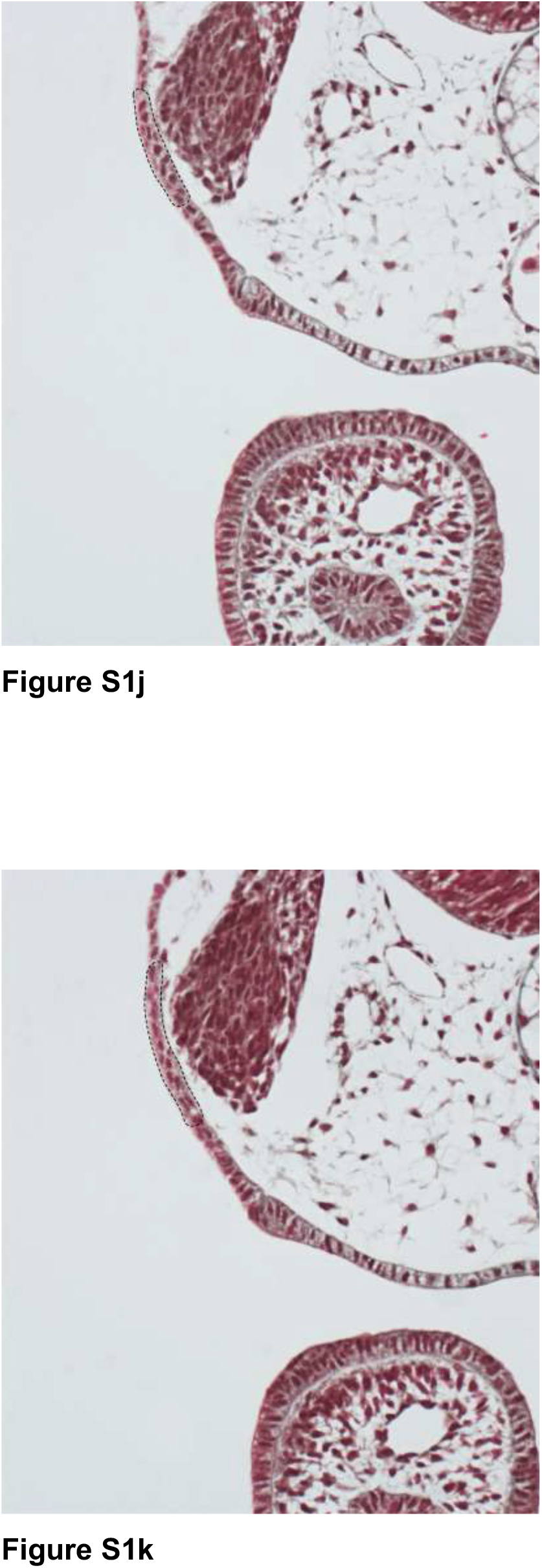
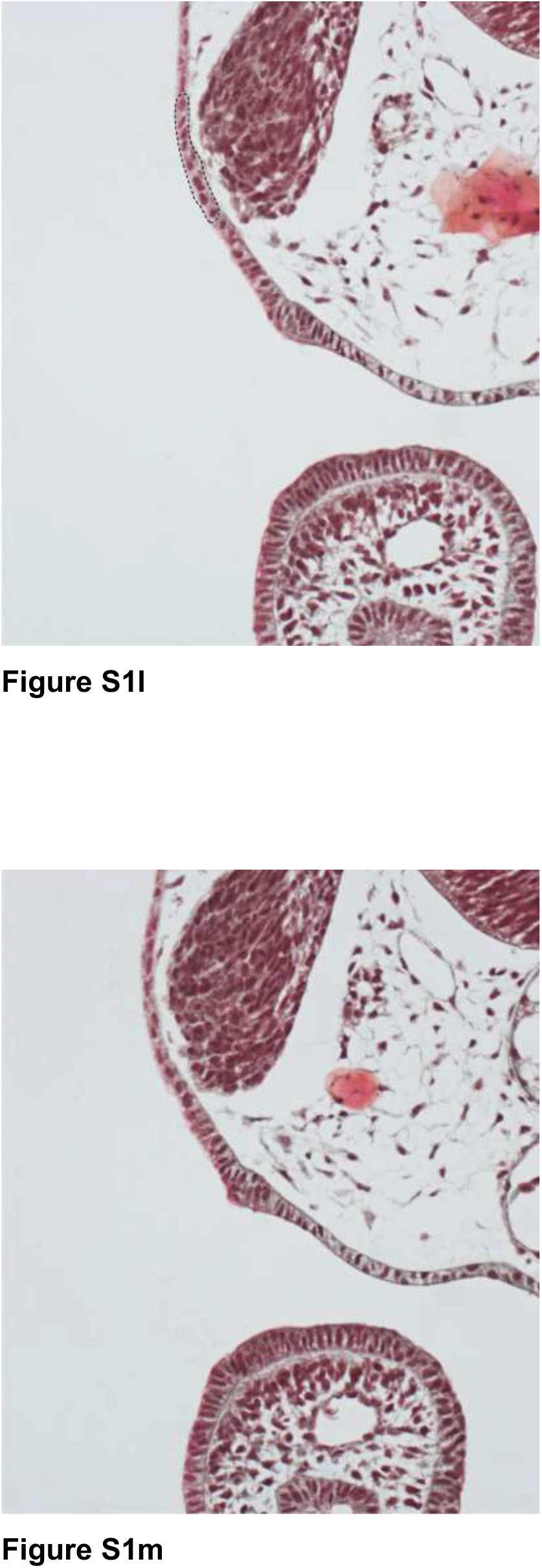
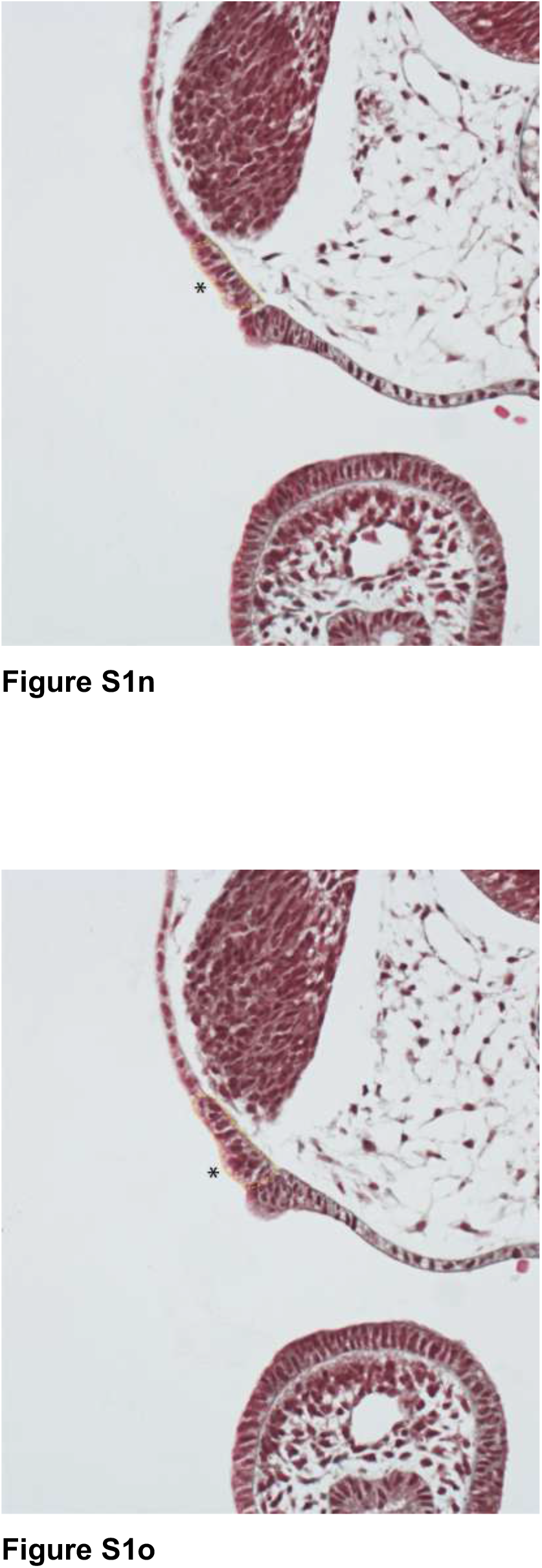
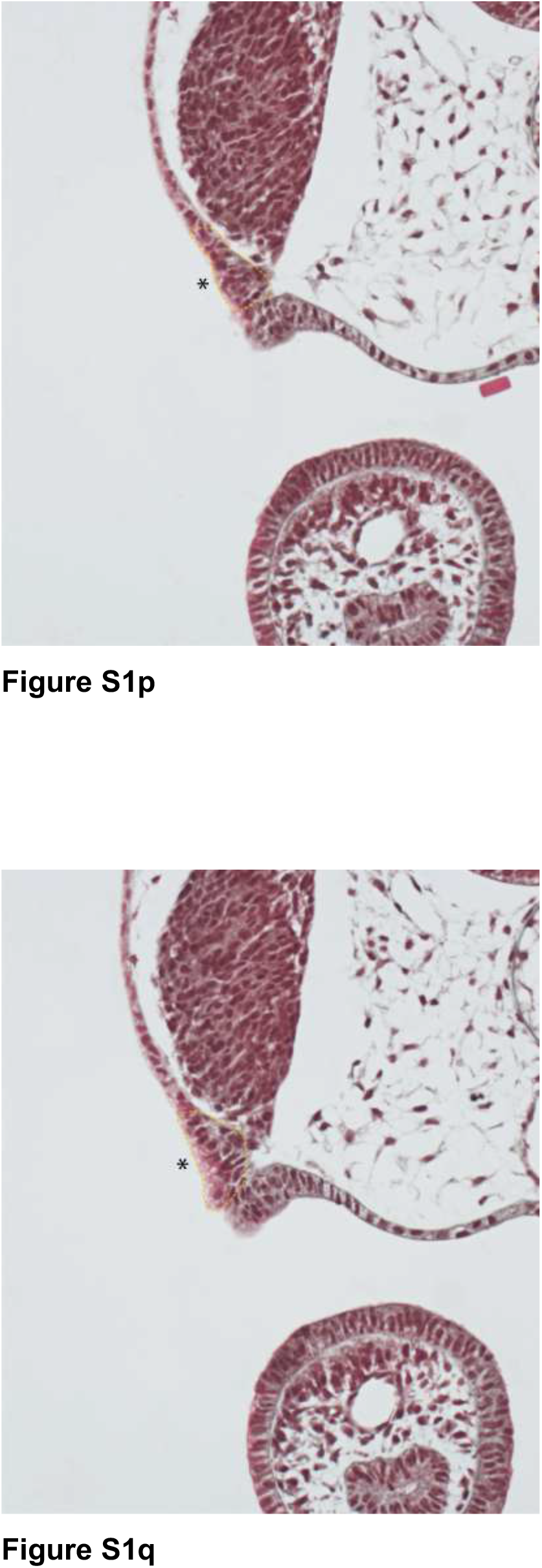
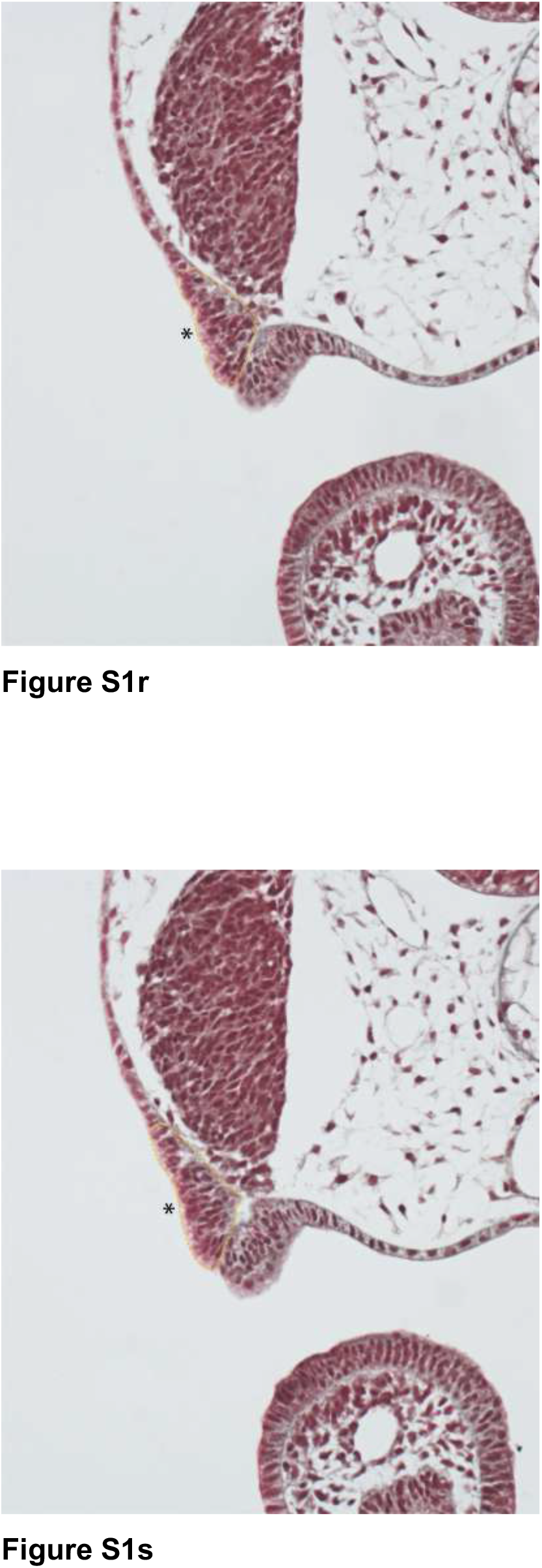
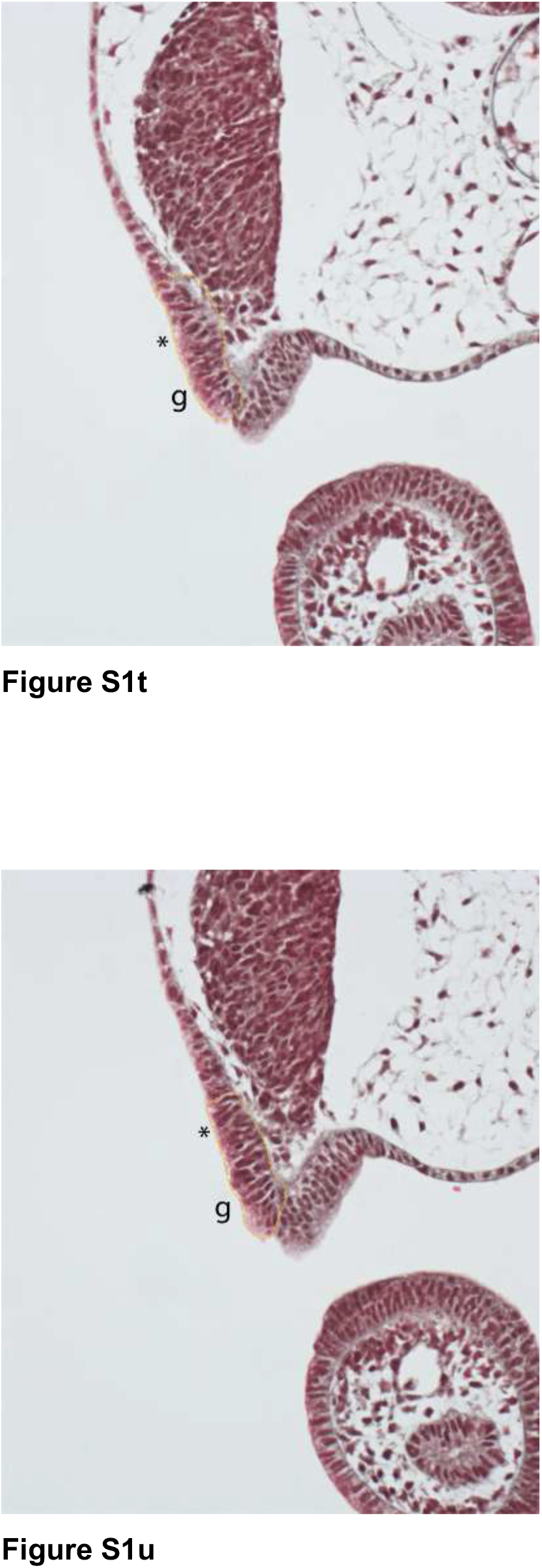
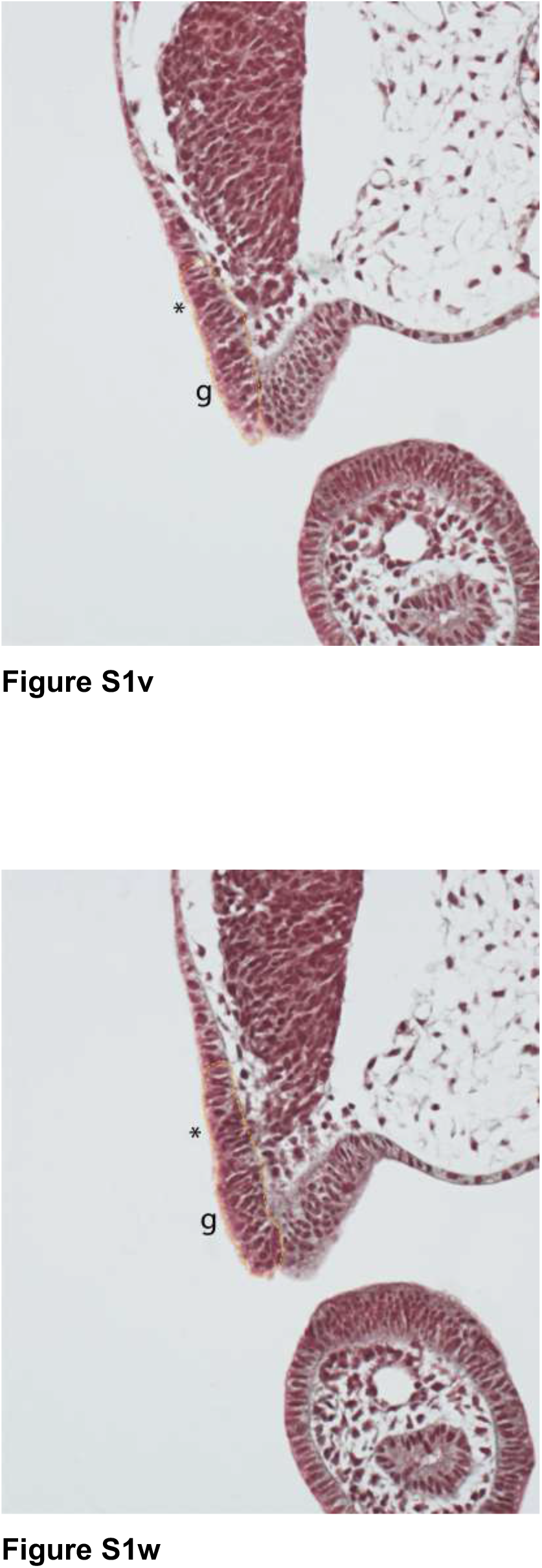
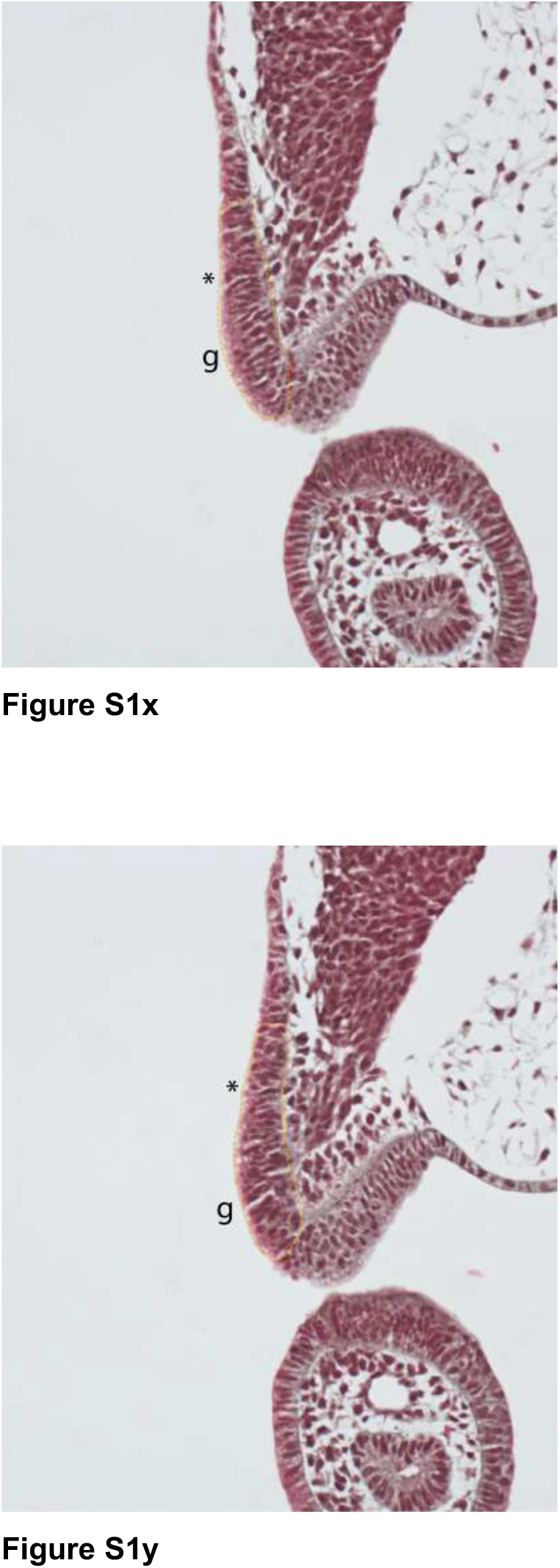
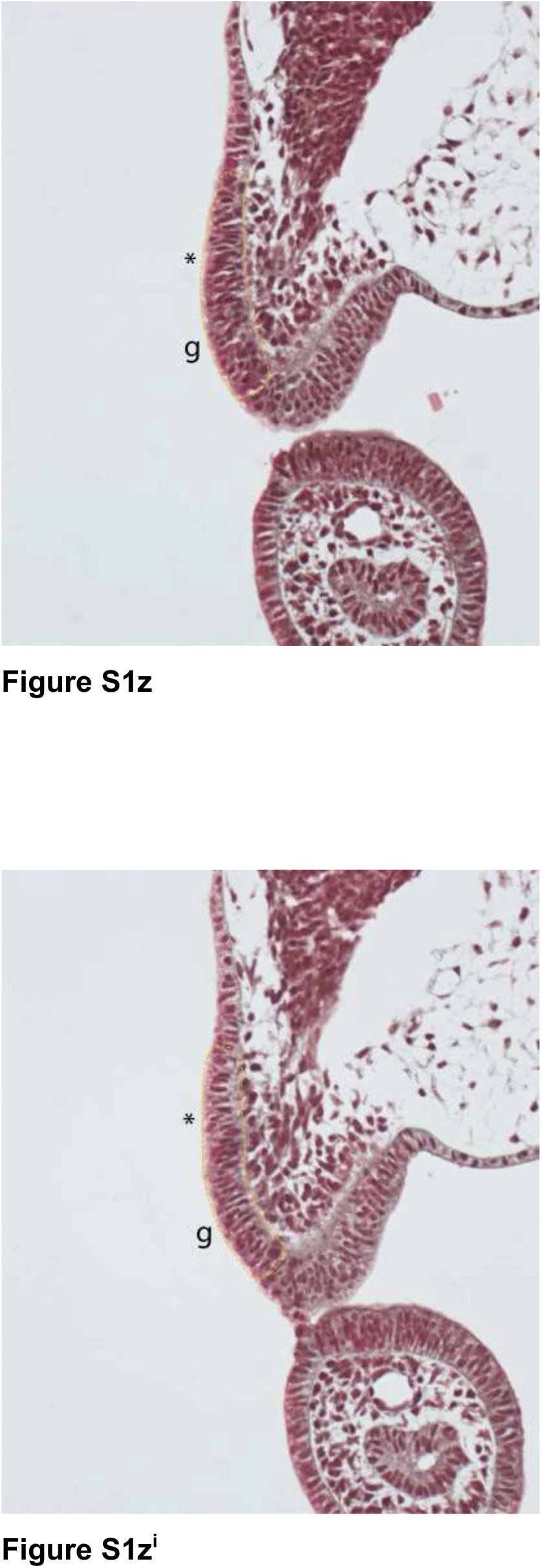
The putative skate spiracular organ placode in relation to the nearest lateral line placode and the geniculate placode. **a-z**^**i**^) A series of adjacent transverse histological sections, in a rostral-to-caudal sequence, at the level of the second (hyoid) pharyngeal arch at S24, illustrating the location of the putative spiracular organ placode (*asterisk*) immediately dorsal to the geniculate placode (*g*), and its distinction from a more dorsal lateral line placode, all giving rise to neuroblasts within a composite ganglion.

## Notes

### Competing Interest Statement

The authors have declared no competing interest.

